# gapTrick – Structural characterisation of protein-protein interactions using AlphaFold

**DOI:** 10.1101/2025.01.31.635911

**Authors:** Grzegorz Chojnowski

## Abstract

The structural characterisation of protein-protein interactions is a key step in understanding the functions of living cells. Here I show that AlphaFold3 often fails to predict protein complexes that are either weak, or dependent on the presence of a cofactor that is not included in a prediction. To address this problem, I developed gapTrick, an AlphaFold2-based approach that uses multimeric templates to improve prediction reliability. I show that gapTrick improves predictions of weak and incomplete complexes based on low-accuracy templates, such as separate protein models rigid-body fitted into a cryo-EM reconstruction. I also show that it identifies with very high precision residue-residue interactions, which are critical for complex formation and a very strong indicator of model correctness. The approach can aid in the interpretation of challenging experimental structures and the computational identification of protein-protein interactions.

**Availability and implementation:** The gapTrick source code is available at https://github.com/gchojnowski/gapTrick and requires only a standard AlphaFold2 installation to run. The repository also provides a Colab notebook that can be used to run gapTrick without installing it on the user’s computer.

## Introduction

Structural characterisation of protein-protein interactions (PPIs) is essential for elucidating roles of protein-coding genes in biological pathways. It’s critical for understanding protein interaction mechanisms, regulatory roles of post-translational modifications (PTMs), and disruptive effects of disease mutations. Experimental approaches like pull-down assays and cross-linking mass spectrometry (XL-MS) can provide some insight into PPI patterns but are difficult to apply for large-scale analyses and often produce inconclusive results ^1^.

Macromolecular crystallography (MX) provides one of the most detailed experimental information on protein structures and was used to determine the majority of structural models available in Protein Data Bank (PDB)^2^. While the crystallisation of small molecules tends to result in optimal packing, macromolecules crystallise in space groups that promote connectivity, preserving internal symmetry of stable complexes ^3^. Therefore, crystallisation can be seen as a natural docking experiment in which multiple alternative interactions between macromolecules form a crystal lattice ^4^. In practice, however, separation of stable, biologically relevant interactions from crystal contacts is not trivial ^5^.

Cryo-EM models are generally easier to interpret because they correspond to complexes that are stable in solution, but similarly to MX they frequently represent small parts of functional complexes in a way that hinders identification of all interactions relevant to their function ^6^. Moreover, cryo-EM models often lack the details needed to reliably identify interactions such as hydrogen bonds or salt bridges, which is crucial for their interpretation and experimental validation. As of January 2025 40% of the EM reconstructions deposited to EMDB ^7^ were determined at a reported resolution below 4Å, meaning that half of their voxels have local resolution significantly worse than 4Å and don’t allow for an accurate model building ^8^.

Artificial intelligence (AI)-based protein structure prediction methods have quickly become an important alternative to the experimental identification of PPIs ^9^. However, they also proved to be computationally expensive and have limited sensitivity. For example, in a recent study targeting binary interactions in a human proteome only 5% of expected interactions were recovered with high confidence ^10^.

The main limitation of the AI-based tools such as AlphaFold2 (AF2) ^11^ is the limited sampling of conformational space near the native state, not their ability to reliably identify a correct prediction ^12^. As a result prediction accuracy of poorly conserved monomeric proteins, with shallow MSAs, strongly depends on the availability of homologous template structures ^11,13^. The contact information derived from templates complements weak evolutionary covariation signals that could be used to identify intramolecular interactions defining a native state of a protein. Although not very well studied, this effect can be even more pronounced for multimeric proteins. Strong protein-protein binding interfaces tend to be highly optimized, involving many interactions that results in a higher evolutionary pressure and sequence conservation ^14^. This may allow for more reliable contact identification and prediction of a native-like structure. Weaker complexes that are less dependent on specific interactions would be more difficult to predict with methods like AlphaFold2. The use of multimeric templates could complement the missing evolutionary information, but it has been deliberately blocked in all AlphaFold versions by masking inter-chain interactions from input templates. Presumably, this was done to avoid potential input bias from low quality templates.

It has been shown that a modification of AlphaFold2, nicknamed AF_unmasked, which uses interchain contacts from input templates, allows predictions of native structures of difficult protein-protein complexes ^15^. However, this approach has been tested on a relatively small set of 251 heterodimeric complexes released after training the AF2-Multimer ^16^ neural network (NN) models, but not otherwise checked for overlap with the training set ^17^. The method can also be computationally expensive in difficult cases.

Here I present gapTrick, an approach that achieves results comparable to AF_unmasked at an order of magnitude lower computational cost. I show that gapTrick can successfully predict PPI structures that cannot be predicted by standard approaches when a suitable multimeric template is given as input. It uses AlphaFold2 NN models trained on monomeric structures allowing validation using any known structure of protein-protein complexes, as none of them have been used for training. Here, I have built a benchmark set of 3,978 stable homo- and hetero-dimeric, trimeric, and tetrameric protein complexes derived from PDB-deposited crystal structures using PISA software ^5^. I show that gapTrick can complement AlphaFold3 (AF3) ^18^ in predicting weak complexes when a template is available, e.g. from a rigid body fit of complex components in a cryo-EM map. I also show that the approach can reliably predict inter-chain interactions in protein-protein complexes. This can help in the interpretation of medium and low-resolution MX and cryo-EM models that don’t allow for a direct identification of residue-residue interactions. The predicted contacts are also a very strong indicator of the model correctness. The approach can help to rebuild experimental models observed in a conformation different from that predicted by AlphaFold2, which otherwise would require tedious rebuilding using interactive software.

The method is nicknamed the “gapTrick” after the original “trick” of combining multiple protein sequences, interspersed with “gaps”, into a single chain to enable prediction of protein complexes with the monomeric AlphaFold2 NN model. It has been discussed on social media ^16^ and implemented by several groups prior the release of AlphaFold2-multimer ^19-21^. None of these implementations combine the trick with the use of multimeric templates.

## Results

### The accurate prediction of weak complexes often requires a template

The PISA software performs an exhaustive analysis of inter-chain contacts in crystal structures to identify subsets of interacting molecules that are potentially stable in solution and may have biological relevance. PISA estimates Gibbs free dissociation energy of a complex (ΔG_diss_) using chemical thermodynamics approximations based on the interface hydrophobicity and specific interactions like hydrogen bonds, salt bridges, disulfide and covalent bonds. In principle, all complexes with ΔG_diss_>0 are stable in solution, but the prediction accuracy increases with the binding energy; it is very high for strong binders but can be overestimated for weaker ones. It has been shown that as a rule of thumb complexes for which the dissociation energy of protein-only components exceeds 40 kcal/mol are stable in over 99% of cases ^4^.

I observed that after removing all non-protein components from the benchmark set structures, the dissociation energy estimate of some of them became negative, clearly showing that their stability depends on the presence of additional components (Figure 1a). AlphaFold3 fails to predict the structure of 38% of complexes with dissociation energy below 50 kcal/mol (TM-score<0.8), but correctly assigns most of them low ranking scores ^18^ below 0.5. The structure of most of these complexes can be correctly predicted with gapTrick if a template (reference model with randomly shifted and rotated complex components) is given on input (Figure 1b). The use of multimeric templates increases the fraction of correct predictions (TM-score>0.8) for targets with dissociation energy below 50 kcal/mol from 62% to 82%.

**Figure 1.**
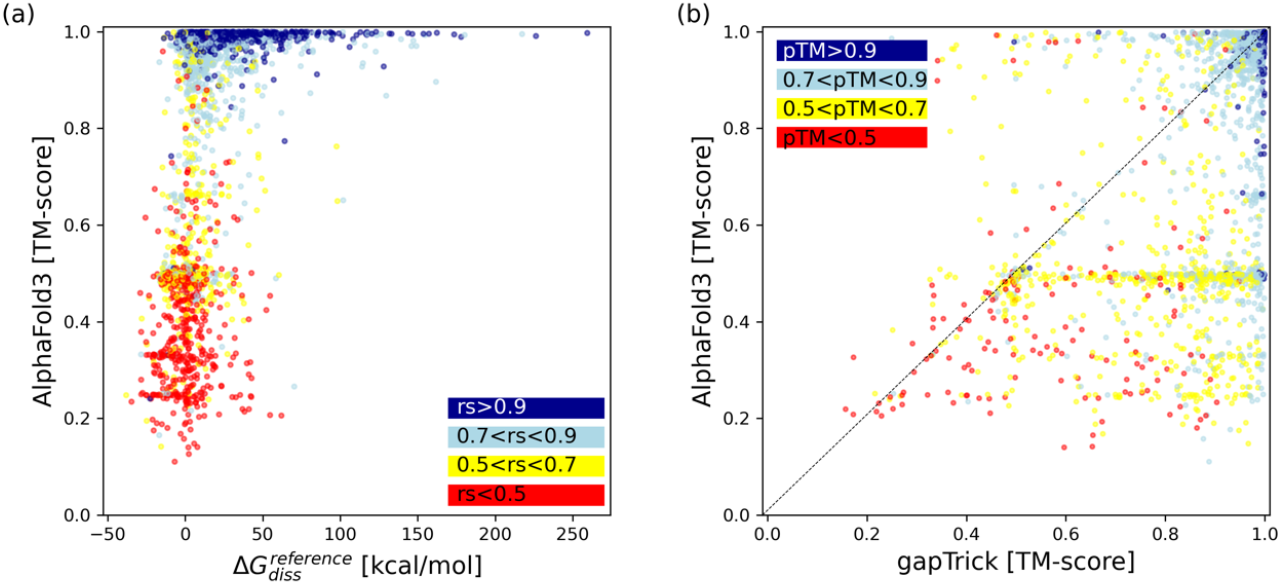
Comparison of AlphaFold3 and gapTrick performance for a benchmark set of dimeric, trimeric and tetrameric protein complex targets predicted by PISA to be stable in solution. The Gibbs free dissociation energy (ΔG_diss_^reference^) was estimated for the reference complexes after removal of all non-protein components. The accuracy of the AlphaFold3 models is significantly reduced for weak protein complexes (a) but can be largely compensated using multimeric templates (b). The colour code indicates the ranking score (rs) of AlphaFold3 (panel a) and the pTM of gapTrick predictions (panel b).

### gapTrick PPI interface contact predictions are highly specific

Reliable identification of residue-residue interactions in protein complexes is important for understanding their function. It’s also a key step of the model validation, which typically relies on structure-driven identification of interactions that are critical for their function *in vivo*. This can be challenging for models, either predicted or experimental, that lack fine structural detail. A natural application for the gapTrick would be rebuilding low-quality, initial models e.g. rigid-body fitted into cryo-EM maps, to identify crucial interactions and help in model interpretation.

Benchmark results show that contacts identified using gapTrick on protein complex interfaces are very precise, even in low recall cases when only a fraction of expected contacts is predicted (Figure 2a). Given the high precision of the contact predictions, a good template, close to a reference structure, will have many contacts that are also predicted by gapTrick with high probability. Indeed, the contact prediction recall strongly depends on the fraction of predicted contacts that are also observed in a template (Figure 2b).

**Figure 2.**
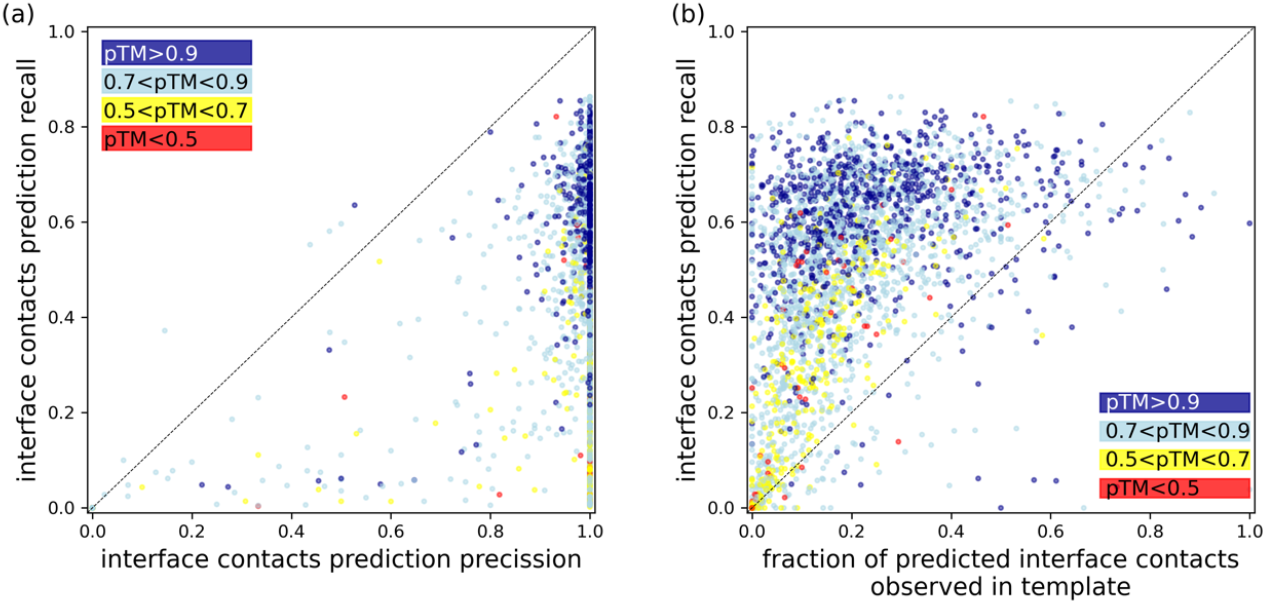
Reliability of the protein-protein interface contacts predicted by gapTrick. The figure shows the recall of inter-chain (interface) contact predictions as a function of contact prediction precision (a) and the fraction of predicted contacts that are satisfied in a template (b). The colour code indicates the pTM score of the gapTrick predictions.

Contact prediction recall for very high confidence models (pTM>0.9) is weakly dependent on the fraction of predicted contacts observed in a template (Spearman’s r=0.34±0.04), implying that they rely largely on information available in MSAs (Figure 2b). However, the recall of interface contact predictions in lower confidence models (pTM<0.9) is significantly reduced for poor templates (Spearman’s r=0.84±0.01), indicating their important role in complementing the weak evolutionary information available in MSAs. The observed high precision of contact predictions agrees with an previous results showing that contact predictions are generally more reliable than experimental coordinates, even though the underlying AlphaFold2 models were trained on them ^22^.

### gapTrick predicts reliable PPI interface coordinates

Although gapTrick predictions provide reliable information on potential residue-level contacts at the complex interfaces, it is not clear how reliable the model coordinates are at the PPI interface, which may be important for model interpretation. To test this, I compared the PISA software estimates of Gibbs free energy of dissociation (ΔG_diss_) for gapTrick predictions and corresponding reference models. PISA is commonly used as a reference for oligomeric-state estimates in PDB-deposited crystal structure models. The program, however, is known to strongly rely on the accuracy of atomic coordinates of the input models, which is required to correctly estimate specific interactions between interface-residues; hydrogen bonds, salt-bridges, and disulfide bonds ^4^.

In the presented benchmarks the Gibbs free energy of dissociation (ΔG_diss_) estimates for the gapTrick models are clearly lower than reference for the small fraction of the predictions (Figure 3a). This is not surprising, as the accuracy of AF2 prediction coordinates has been estimated to be comparable with medium to low resolution crystal structures ^23^. The issue, however, affects mainly low-confidence models (Figure 3b). Although there is a strong linear correlation between the two dissociation energy estimates (Pearson’s r=0.90±0.01), it is noticeably lower for medium to low confidence predictions with pTM<0.7 (Pearson’s r=0.58±0.04). There is also a small, negative linear correlation between the dissociation energies difference between predicted and reference models and pTM (Pearson’s r=-0.28±0.02), indicating that ΔG_diss_ is often underestimated for low confidence predictions.

**Figure 3.**
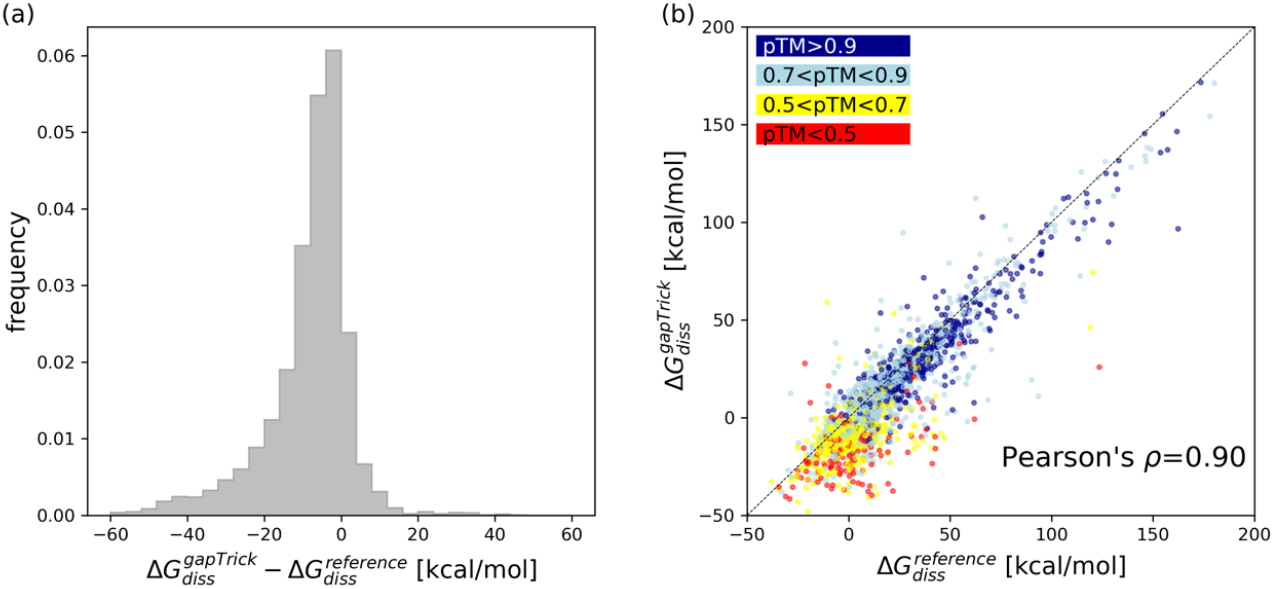
Comparison of Gibbs free energy of dissociation estimates for gapTrick predictions (ΔG_diss_^gapTrick^) and reference models (ΔG_diss_^reference^). The distribution of a difference between gapTrick and reference model ΔG_diss_ (a) and a scatterplot showing their strong linear correlation (b). The dashed line in panel (b) shows the diagonal of the plot and a perfect correlation of the abscissa and ordinate values. The colour code indicates the pTM score of the gapTrick predictions.

### Identification of correct complexes based on gapTrick contact predictions

So far, I only considered structurally characterised protein-protein complexes with good evidence that they form stable interactions. For some gapTrick applications, however, it would be very important to verify the correctness of predicted models, for example in cases where several alternative templates exist.

To test this, I created three benchmark sets consisting of AlphaFold3 predictions for pairs of proteins unlikely to form complexes (hu.MAP2 negative PPIs), likely to form complexes (hu.MAP2 positive PPIs), and models extracted from PDB deposited cryo-EM structures resolved at a resolution worse than 4Å.

In the absence of reference models, it is not possible to evaluate the accuracy of AF3 predictions for hu.MAP2 positive/negative PPIs. However, as the AF3 ranking scores are significantly higher in the positive test (with a t-test p-value below 0.0001), it can be assumed that at least some of the structures in the positive set are correct. Furthermore, due to AF3’s tendency to hallucinate ^18^, it can be expected that most of the models will look plausible and have good stereochemical properties even if they are completely wrong.

All AF3 predictions and cryo-EM models were automatically rebuilt using gapTrick. The difference in the number of predicted protein-protein interface contacts is a very strong discriminator between the sets (Figure 4a). It can also be used to construct a reliable classifier that discriminates between very unlikely but geometrically plausible (negative PPIs) and structurally validated cryo-EM complexes (Figure 4b). It has much stronger predictive power than the pTM scores estimated by gapTrick (monomeric AF2 NN models don’t estimate ipTM, Figure 4b). This clearly demonstrates that the number of predicted contacts at the protein-protein interface can be used to identify plausible predictions. According to the benchmarks, models with as few as 3 predicted contacts should be correct in over 85% of cases, with a false positive rate of 16%, which corresponds to precision and recall of 86% and 85% respectively. This is much higher than the classification performance obtained by Humphreys et al ^21^ using monomeric AF2 NN models without templates, based on the probability of a single, most-likely inter-chain contact below 12Å.

**Figure 4.**
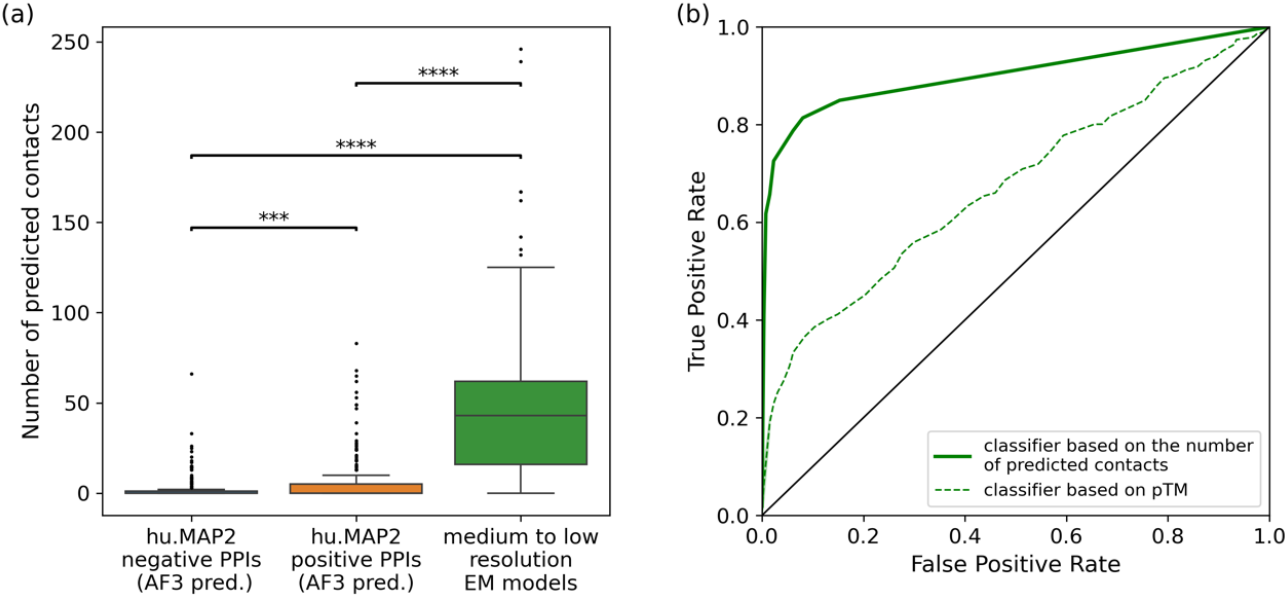
(a) Comparison of distributions of the number of protein-protein interface contacts predicted by gapTrick based on templates from AF3 complex predictions for hu.Map 2.0 negative and positive PPI sets or medium to low resolution cryo-EM binary complexes extracted from PDB. Stars indicate significance of two-sample t-tests with the hypothesis that the expectation value of a distribution on the left is smaller. (b) ROC curves of classifiers based on pTM or the number of protein-protein interface contacts predicted by gapTrick (area under the ROC curve 0.67 and 0.90).

### Characterisation of protein-protein interactions in cryo-EM maps

Phospholipase Cγ (PLCγ) enzymes control many vital cellular processes and their mutations can lead to abnormal cell signalling and the development of cancer. In a recent study, a complex of the human hPLCγ2 with the regulatory tyrosine kinase FGFR1 was determined using cryo-EM ^24^, but the 4.1Å resolution of the reconstruction didn’t allow for a mechanistic description of the complex formation (PDEB/EMDB ids 8jqi/36573, Figure 5a). Although a highly conserved, phosphorylated Tyr766 of FGFR1 was previously identified as the major site of interaction between FGFR1 and PLCγ2, all the structural context was available from a related crystal structure of an FGFR1 in complex with rat rPLCγ1 (PDB id 3gqi)^25^. Here, I show that gapTrick can successfully provide the structural context of the PLCγ2-FGFR1 complex formation based on the medium resolution cryo-EM map alone.

**Figure 5.**
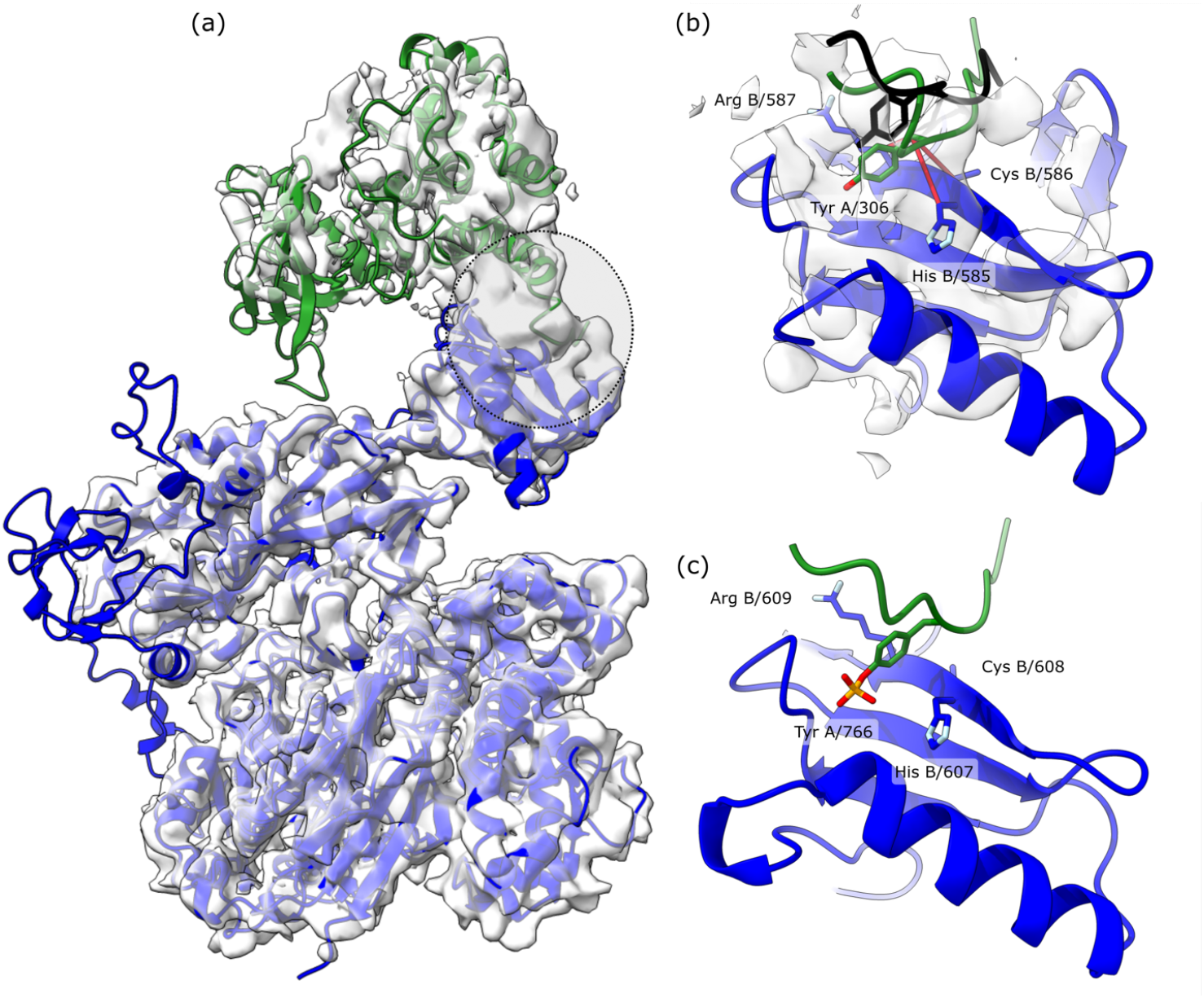
Cryo-EM structure of a human phospholipase hPLCγ2 in complex with tyrosine kinase FGFR1 resolved at 4.0Å resolution (a). Panel (b) shows a noisy map region at the interface of the hPLCγ2-FGFR1 complex, and the corresponding fragment of the model built fitted to the map using gapTrick. The region is highlighted with a dashed circle on panel (a). Red lines indicate high probability contact predictions between hPLCγ2 interface residues and a highly conserved Tyr766 of FGFR1. A fragment of the deposited FGFR1 model (PDB 8jqi) superposed by the displayed fragment of the hPLCγ2 (residues A/530-630, Cα rmsd 1.4Å) is shown in black. Panel (c) shows a corresponding fragment of a 2.5Å resolution crystal structure model of a related complex of human FGFR1 with rat rPLCγ1 (PDB 3gqi). The residues predicted by gapTrick to be in contact in the hPLCγ2-FGFR1 complex are strictly conserved in rPLCγ1 and have been identified as part of a canonical FGFR1 binding site. Phospholipases rPLCγ1 and hPLCγ2 are shown in blue, FGFR1 in green.

AlphaFold3 fails to predict the structure of the complex, with and without phosphorylation of Tyr766 (ipTM/TM-score 0.35/0.49 and 0.55/0.49, respectively). Therefore, I built the model from AF2 structure predictions of individual chains (AlphaFold Protein Structure Database ^26^ version 4 models for Uniprot P16885 and P11362). The chains were fitted into the map using MOLREP^27^, refined in real space with self-restraints generated at the 4.3Å cutoff in COOT^28^, and rebuilt using gapTrick. The resulting prediction had a pTM of 0.81 and excellent stereochemical properties (MolProbity score of 1.15). It required only 20 cycles of automated refinement using Servalcat^29^ with jelly body restraints and default parameters; no interactive rebuilding was required. The resulting model has map-model fit parameters comparable to the deposited model (CC_mask_ of 0.73 and 0.72, for deposited and rebuilt models, respectively). Although no rotamer or Ramachandran plot restraints were used during refinement, the final model has very good stereochemical properties (MolProbity score 1.23 compared to 2.25 for the deposited model).

The prediction includes several high probability contacts at the hPLCγ2-FGFR1 interface (Figure 5b). The contacts correspond to the previously identified canonical binding site of the phosphorylated Tyr766 of FGFR1 and closely resemble a previous crystal structure model of human FGFR1 with a rat phospholipase rPLCγ1 (Figure 5c) ^25^. The Gibbs free energy of dissociation estimated for the new model is -1.5 kcal/mol and comparable to the deposited hPLCγ2-FGFR1model (0.5 kcal/mol) and the corresponding rPLCγ2-FGFR1 crystal structure (1.8 kcal/mol). In the light of the results presented in this paper, this relatively weak complex would be difficult to predict, regardless of the well-studied and biologically relevant interaction mechanism.

### Comparison between gapTrick and AF_unmasked

The use of multimeric templates was deliberately disabled by the AF2 developers. However, a recent publication has shown that the software can be modified to allow the use of inter-chain contact information from templates to improve prediction performance with AF_unmasked^15^. In addition to an artificially generated benchmark set, the AF_unmasked method was tested on a set of 28 heterodimeric protein complexes that cannot be predicted by AlphaFold2 (DockQ<0.2), but for which a homologous complex is available in the PDB (DockQ>0.15). The authors showed that most of the test set structures can be correctly predicted based on the homologous templates.

Here, I compared performances of gapTrick and AF_unmasked on this test set. For comparison, I used predicted structures provided by the AF_unmasked authors to avoid any issues related to a non-optimal configuration of their software. The gapTrick predictions were generated fully automatically using default parameters. The model accuracies (TM-score against a reference) of the two approaches are highly correlated (Figure 6). They are also significantly better than the AF3 predictions, although it is not clear whether the set of heterodimeric complexes and the AF3 training set share any homologous structures that might bias the predictions.

**Figure 6.**
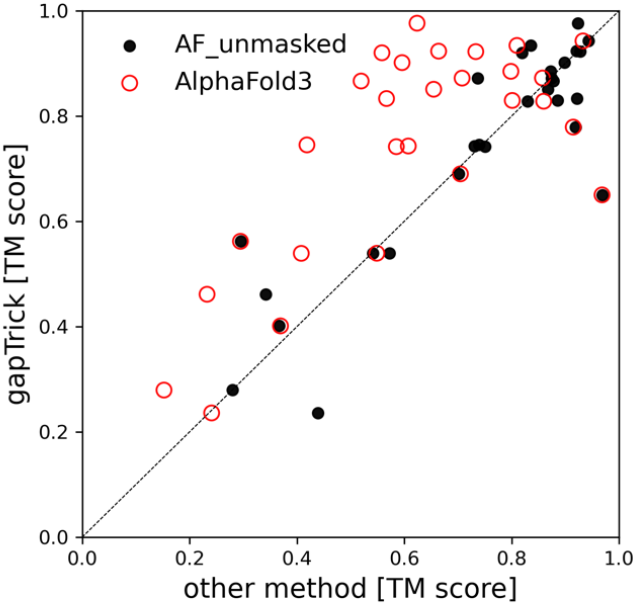
Comparison of the performance of gapTrick, AF_unmasked and AlphaFold3 on a set of 28 heterodimeric proteins defined by Mirabello et al ^15^. Both gapTrick and AF_unmasked predictions were generated using complexes homologous to the target as templates.

A more comprehensive comparison of gapTrick and AF_unmasked is beyond the scope of this work due to the problems with defining the benchmark strategy, which is much more difficult for the latter. Most importantly, it is not trivial to avoid information leakage between training and test sets for a model trained on protein complexes^17^. Furthermore, unlike gapTrick, AF_unmasked requires user-defined mapping between template chains and target sequences and is not applicable for automated, large-scale processing of higher oligomers. Finally, only gapTrick provides tools for the analysis of predicted contacts.

## Discussion

I show that AlphaFold3 often fails to predict structures of weak complexes whose formation may be transient or dependent on the presence of additional components, which are not included in the prediction. The prediction of such complexes is crucial for the identification of novel PPIs for which a complete composition, required for a stable complex formation, has not yet been identified. Modelling of small fragments of protein complexes may be also important for building very large cryo-EM models ^30^.

Many weak complexes can be predicted using gapTrick when a low accuracy multimeric template is already available. The templates can be homologous structures, or rigid-body fits of the complex components to a cryo-EM map using approaches like SliceNDice ^31^. Templates can also originate from *de novo* cryo-EM map interpretation approaches such as ModelAngelo ^32^, which typically produce fragmented and incomplete models in regions of lower local resolution. In such cases, gapTrick can support tedious and error-prone model building using interactive methods. It can also complement automated approaches such as PDB_REDO ^33^ or CERES ^34^. Particularly important here are excellent stereochemical properties of models predicted by gapTrick (Figure S1). The model geometry should not deviate from energetically favourable conformations unless there is very good evidence in the data supporting unusual conformations ^35^.

Models of protein complexes, like any other scientific model, require experimental validation, which typically relies on structure-driven identification of interactions that are critical for their function *in vivo*. I have shown that gapTrick can identify crucial contacts in low-accuracy models of protein complexes whose overall structures cannot be accurately predicted using standard approaches. The contacts are predicted with high precision and can help to assess model correctness and to define residues for mutagenesis in a functional analysis. They are also a strong indicator of model correctness, which will be very important in cases where there is no other evidence of complex formation and topology.

## Acknowledgements

The manuscript was written without the use of AI, but revised using DeepL Write to address issues related to the fact that I am not a native English speaker. Like AI, non-artificial intelligence never operates in isolation and gains knowledge from its environment. Therefore, I would like to thank the authors of the referenced publications for sharing their thoughts, results, and code, which were an invaluable help in obtaining presented results. I would like to thank Matthias Wilmanns, Jan Kosiński, Dima Molodenskiy, and colleagues from EMBL, CCP4, and CCP-EM for fruitful discussions. I thank Daniel J. Rigden and Eugene Krissinel for critical reading of the manuscript and very helpful comments. Finally, I would like to thank the EMBL IT for providing the HPC computational resources.

## Methods

For simplicity, I use the following assumptions and definitions throughout the paper:

- Two residues are in contact if the distance between their Cβ atoms (Cα for glycine) is less than 8Å. This does not imply a physical interaction (e.g. a hydrogen bond formation)
- The gapTrick contact predictions are based on AlphaFold2 distograms^11^. Here, only inter-chain contacts with probability greater than 0.8 are analysed
- All gapTrick benchmarks are based on templates generated from reference models with each chain rotated and translated by random angles and distances, as described in Materials and methods

### Code availability

The gapTrick source code is available at https://github.com/gchojnowski/gapTrick and requires only a standard AlphaFold2 installation to run. The repository also provides a Colab notebook that can be used to run gapTrick without installing it on the user’s computer.

### Description of the gapTrick algorithm

The prediction of protein-protein complexes in gapTrick is based on AlphaFold2 NN models trained on single-chain proteins. The method uses only two out of five AF2 NN models that allow for the use of input templates (1 and 3). Therefore, by default, only two structural models are generated for each target, compared to 25 in AlphaFold-Multimer and AF_unmasked (5 predictions for each of the 5 NN models). To allow prediction of multimeric structural models, all input protein chains are merged into a single protein chain interspersed with 200 amino acid gaps. The only input required from user is a PDB/mmCIF template and a FASTA file containing all target sequences in arbitrary order. The structure prediction is performed fully automatically in the following steps:

1. Multiple Sequence Alignments (MSAs) are generated separately for all input chains. This step uses the MMseqs2 API^36^ by default, but the alignments can be also provided by the user in a3m format.
2. Target sequences are assigned to the template chains using a greedy algorithm maximising sequence identity
3. Following the sequence assignment the template chains and target sequences are merged into single protein chains with residue index gaps (200 by default)
4. MSAs for successive target sequences are merged into a single MSA. No sequence pairing is used at this step, as it’s not required for successful prediction of protein complexes^37^
5. The resulting single-chain protein structure is predicted using a standard AlphaFold2 pipeline with default parameters
6. The predicted models are broken down into multiple chains and ranked by the pTM score (ipTM score is not predicted by monomeric AF2 NN models). Top-ranked model is subjected to the AMBER force field relaxation.

### Benchmark set of MX structures from PDB

A benchmark set of stable in solution protein complex structures was generated using PISA software based on PDB deposited protein-only crystal structures. Since the reliability of the PISA analysis depends heavily on the quality of the model coordinates^4^, for the analysis I selected protein-only crystal structure models deposited with experimental data, solved at a resolution of 2.5Å or better and with R_free_ less than 0.25. For computational efficiency of model-prediction, I additionally restricted the set to structures with molecular weight between 20 and 50 kDa. As of 23.09.2024 this procedure resulted in 13,539 protein crystal structure models that were subsequently processed by PISA to identify stable dimeric, trimeric, and tetrameric protein complexes (both homo and hetero oligomers were accepted). From each crystal structure model only a single top-ranked complex of a desired multimeric state with all protein chains longer than 50 amino acids was selected. All selected complexes have a non-negative Gibbs free dissociation energy (ΔG_diss_). However, after removing non-protein components from the structures (ions, waters, and small molecules), the energy estimate for some of the complexes became negative, suggesting that the presence of the removed components may be critical for their stability. These structures have been retained in the benchmark set as they reflect realistic scenarios for modelling complexes before components critical to their stability have been characterized. The final benchmark set contains 3,978 structural models of protein-protein complexes, including 2,000 dimers, 722 trimers, and 1,256 tetramers.

To mimic low-accuracy experimental models (e.g. rigid-body fitted in cryo-EM maps), templates for the use with gapTrick were generated from reference models by rotating and translating all chains about their centres of mass by uniformly distributed random values up to 30 degrees and 5 Å. This procedure is hard-coded into gapTrick and can be invoked with the *--rotrans=30,5* keyword.

It is not a purpose of this work to systematically benchmark AlphaFold3, but to show that its performance may be improved for some cases, using additional data. Therefore, the benchmark set used here completely neglects any potential overlap with the AF3 training set.

### Benchmark set of putative binary complexes

From the PPIs defined by hu.MAP 2.0 database^38^, a positive and negative sets of 300 binary complexes each were randomly selected and their structures predicted using AlphaFold3 with default parameters.

Experimentally characterised binary complexes were extracted from PDB-deposited cryo-EM structures determined at resolutions between 4Å and 10Å using PISA. From all potential complexes identified by the program, only those with interface hydrophobicity p-value below 0.5 were selected. As of 15 February 2025, 320 models of binary complexes were extracted using these criteria.

### Numerical methods

Correlation coefficients shown in the results are bootstrap estimates with 100 repeats. All the statistical analyses were performed using tools implemented in Scipy^39^ and Numpy^40^. Figures were prepared using matplotlib^41^ and ChimeraX^42^. AlphaFold3 predictions were obtained with default parameters using a stand-alone installation of a version 3.0.0 and weights downloaded on 6.12.2024. The gapTrick code is based on AlphaFold version 2.3.2 and uses AF2 NN models dated 14.07.2021. The cryo-EM model was built and refined using CCP-EM v2^43^, MOLREP version 11.9.020.9^27^, Servalcat version 0.4.32^29^, and COOT version 0.9.8.93^28^. For model validation and analysis I used MolProbity^44,45^ and PISA distributed with CCP4 version 8.0.018^46^. Molecular models processing was performed using scripts based on Gemmi^47^, CCTBX^48^, and BioPython^49^. Multiple sequence alignments used for benchmarks were prepared using a standalone version of MMseqs2 release 14^36^ and a colabfold_search script^20^. The gapTrick release by default uses MMseqs2 API. Both standalone MMseqs2 installation and API were invoked with *--use-env 0* and *--filter 0* options. The TM-scores of predicted structures to the reference models were calculated using Foldseek release 9^50^ using the easy-multimersearch option.

## Supplement: Stereochemical properties of models rebuilt with gapTrick

A convenient application of gapTrick is rebuilding and completion of models fitted into cryo-EM or MX maps. It can be particularly useful when a high-confidence prediction of a complete complex is not available and low map resolution doesn’t allow *de novo* model tracing. In such cases, the complex models must be assembled in the maps from individual chains, which doesn’t allow for detailed modelling of PPI interfaces. I have shown that gapTrick can successfully remodel such structures and predict contacts that may be crucial for assessing their correctness. However, the overall plausibility of the stereochemical properties of such models remains an open question.

The stereochemical properties of macromolecules have been studied in detail over the years using both theoretical and statistical approaches to define energetically favourable conformations. The model geometry should not deviate from these values unless there is very good evidence in the data supporting an unusual conformation, which is usually not the case at lower resolutions ^35^. Therefore, reliable stereochemical properties of gapTrick predictions are crucial, as they facilitate refinement, model interpretation, and subsequent deposition to the PDB.

**Figure S1.**
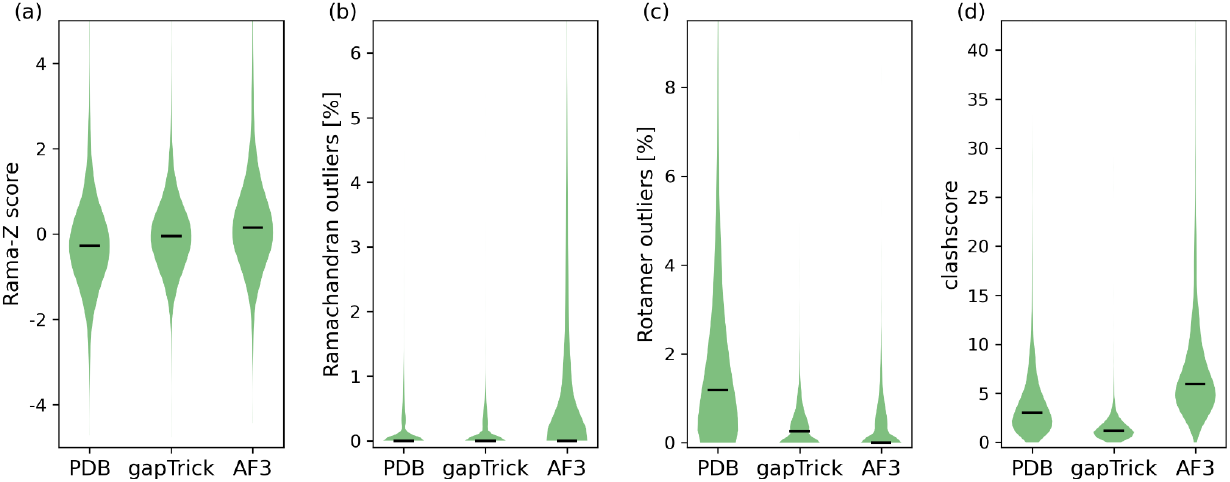
Comparison of the most important MolProbity validation scores estimated for models predicted using gapTrick, AlphaFold3 (AF3), and PDB-deposited models. To reduce bias due to the presence of experimentally unresolved, and often intrinsically disordered regions, the predicted models were truncated to residue ranges covered by PDB-deposited structures prior to scoring. Horizontal lines on the violin plots represent medians.

The models predicted by gapTrick have stereochemical properties reported by MolProbity^44^ that are comparable to or better than those of the PDB-deposited crystal structure models and AlphaFold3 predictions (Figure S1). They have reliable distributions of backbone torsion angles (Figure S1a), with only 1% significantly different from those observed for high quality crystal structures (absolute value of Rama-Z score above 3)^45^. The corresponding fractions for PDB-deposited and AF3-predicted models from the benchmark set are 2% and 3%, respectively. The gapTrick predictions have a comparable number of Ramachandran plot outliers to the fully refined crystal structure models (Figure S1b). They have very few non-rotameric side chains, which is very important for the interpretation of medium and low-resolution maps (Figure S1c). Finally, the gapTrick models have a very low clashscore (number of steric conflicts per 1000 atoms in the model) with a median of 1.2 (Figure S1d). This is significantly lower than the median clashscore of the input templates, with randomly shifted and rotated chains, which is as high as 44. These results clearly show that gapTrick automatically resolves most of the stereochemical issues in the templates, and that they do not require additional pre-processing and rebuilding.

